# Lignin isolated by microwave-assisted acid-catalyzed solvolysis induced cell death on mammalian tumor cells by modulating apoptotic pathways

**DOI:** 10.1101/2024.01.20.576161

**Authors:** Rio Kashimoto, Eriko Ohgitani, Yutaka Makimura, Tatsuya Miyazaki, Chihiro Kimura, Masaharu Shin-Ya, Hiroshi Nishimura, Giuseppe Pezzotti, Takashi Watanabe, Osam Mazda

## Abstract

Lignin, the most abundant renewable aromatic polymer, has been shown to suppress the growth of mammalian tumor cells. Despite being a textbook example, there is little information on the biological activity of lignin in relation to its molecular structure or the molecular mechanisms by which lignin suppresses tumor cells in mammalian species. Here, we prepared Microwave-assisted Acid-catalyzed Solvolysis Lignin (MASL) and assessed its effects on human and mouse tumor cells. Our data showed MASL significantly reduced viability of tumor cells by modulating apoptotic pathways. MASL treatment upregulated TNF-α, Fas, and FasL expression levels, while suppressing anti-apoptotic NF-κB and mTOR pathways in tumor cells. In-vivo experiments were also performed using tumor-bearing mice to confirm the anti-tumor effects of MASL. An administration of MASL significantly suppressed tumor growth in mice in association with elevation of caspase 3 expression. These findings strongly suggest the potential usefulness of low molecular weight lignin as an effective therapeutic against malignancies.

## 1. Introduction

Lignin, a heterogeneous aromatic polymer, stands as the second most abundant biopolymer, constituting 5-35% of plant cell walls [1]. Lignin enhances plant cell wall rigidity, facilitates mineral and water transport through vascular bundles and provides protection against pests and pathogens. These abilities extend beyond plant species, with previous reports suggesting lignin’s potential to suppress mammalian tumor cells. Lignin nanoparticles have been studied as drug carrier for colon cancer therapy, demonstrating minimal induction of drug resistance [2, 3]. Lignin-carbohydrate complexes (LCCs) derived from pine cones have anti-tumor effects on ascites sarcoma-180 cells [4]. Moreover, lignin-rich enzyme lignin (LREL) activated dendritic cells via receptor protein, TLR4, and enhanced expression of CD86, IL12p40, and TNF-α in dendritic cells [5]. However, the structure-function relationship for the activity has not been clearly elucidated, especially at the cellular process and molecular levels, has not been clearly elucidated [6-11].

To clarify the biological and pharmacological effects of the plant-derived lignin fractions with detailed structural data, we extracted lignin by applying microwave acid-catalyzed solvolysis of softwood and hardwood and subsequent organic solvent extraction [12, 13, 14]. Microwave solvolysis in hydrophobic solvents containing alcohol and acids decomposes lignin selectively to give lignin oligomers and monomers without condensation reactions of lignin units due to trapping of unstable enol ether intermediates by the alcohol [10, 11]. The reaction products retained higher amount of native lignin interunit linkages than those obtained with acidolysis, sulfite cooking, kraft pulping and alkaline treatment under harsh conditions. Extraction of the degradation products with toluene, ethyl acetate and acetone give soluble lignin oligomers with different molecular weight distributions, facilitating application to bioassay [15, 16]. Thus, MASL, lignin molecules with reduced molecular weight and chemical modification expands the potentials of lignin as bioactive agent.

Our experimental data demonstrate that MASL significantly induced anti-tumor effects on mouse and human tumor cells in vitro and in vivo. Furthermore, we determined that tumor cell death caused by MASL may be mediated by modulation of pro- and anti-apoptotic signaling pathways. Our analysis represents a preliminary step in advancing the understanding of lignin treatment in the cell death pathway, elucidated through both in vivo and in vitro experiments on malignant tumor cells.

## 2. Materials and Methods

### 2.1 MASL samples

All reagents were of analytical grade and were purchased from (Wako Pure Chemical Industries Ltd. Osaka, Japan) and (Nacalai Tesque). Eucalyptus globulus and Japanese cedar wood were obtained from (Nippon Paper Industries Co., Ltd.). E. globulus wood particles, Japanese cedar wood particles, and alkaline lignin from Japanese cedar wood were degraded by microwave solvolysis with a mixture (20 mL) of toluene, ethanol, and water (8:6:6) containing 0.75 g or 0.25 g of H_2_SO_4_ using a microwave reactor, (Biotage Initiator Plus) at 180° C for 30 min (Table 1 and Table 2) as described [12, 13, 17]. After reactions, low-molecular-mass lignin was extracted with toluene, ethyl acetate, and acetone (Entry 1-12). The same lignin fractions were prepared by a scale-up process with a mixture (200 mL) of toluene, methanol, and water (6:6:8) containing 7.5g of H_2_SO_4_ using a microwave reactor (Milestone StartSYNTH) at 180° C for 30 min (Entry 13-15). Organic solvent extracts from the scale-up process were also prepared with a mixture (200 mL) of toluene, ethanol and water (6:6:8) containing 7.5 g of H_2_SO_4_ (Entry 16-18).

**Table 1.** MASL used in this study.

| Sample | Resources | Reagents for microwave solvolysis |
| --- | --- | --- |
| YM C1T | Japanese cedar wood particles | H<sub>2</sub>O, EtOH, toluene (6:6:8), 0.75 g H<sub>2</sub>SO<sub>4</sub> |
| YM C1E | Japanese cedar wood particles | H<sub>2</sub>O, EtOH, toluene (6:6:8), 0.75 g H<sub>2</sub>SO<sub>4</sub> |
| YM C1A | Japanese cedar wood particles | H<sub>2</sub>O, EtOH, toluene (6:6:8), 0.75 g H<sub>2</sub>SO<sub>4</sub> |
| YM E1T | *E. globulus* wood particles | H<sub>2</sub>O, EtOH, toluene (6:6:8), 0.75 g H<sub>2</sub>SO<sub>4</sub> |
| YM E1E | *E. globulus* wood particles | H<sub>2</sub>O, EtOH, toluene (6:6:8), 0.75 g H<sub>2</sub>SO<sub>4</sub> |
| YM E1A | *E. globulus* wood particles | H<sub>2</sub>O, EtOH, toluene (6:6:8), 0.25 g H<sub>2</sub>SO<sub>4</sub> |
| YM C2T | Japanese cedar wood particles | H<sub>2</sub>O, EtOH, toluene (6:6:8), 0.25 g H<sub>2</sub>SO<sub>4</sub> |
| YM C2E | Japanese cedar wood particles | H<sub>2</sub>O, EtOH, toluene (6:6:8), 0.25 g H<sub>2</sub>SO<sub>4</sub> |
| YM C2A | Japanese cedar wood particles | H<sub>2</sub>O, EtOH, toluene (6:6:8), 0.25 g H<sub>2</sub>SO<sub>4</sub> |
| YM E2T | *E. globulus* wood particles | H<sub>2</sub>O, EtOH, toluene (6:6:8), 0.25 g H<sub>2</sub>SO<sub>4</sub> |
| YM E2E | *E. globulus* wood particles | H<sub>2</sub>O, EtOH, toluene (6:6:8), 0.25 g H<sub>2</sub>SO<sub>4</sub> |
| YM E2A | *E. globulus* wood particles | H<sub>2</sub>O, EtOH, toluene (6:6:8), 7.5 g H<sub>2</sub>SO<sub>4</sub> |
| YM CL1T | Japanese cedar alkali lignin | H<sub>2</sub>O, MeOH, toluene (6:6:8), 7.5 g H<sub>2</sub>SO<sub>4</sub> |
| YM CL1E | Japanese cedar alkali lignin | H<sub>2</sub>O, MeOH, toluene (6:6:8), 7.5 g H<sub>2</sub>SO<sub>4</sub> |
| YM CL1A | Japanese cedar alkali lignin | H<sub>2</sub>O, MeOH, toluene (6:6:8), 7.5 g H<sub>2</sub>SO<sub>4</sub> |
| YM CL2T | Japanese cedar alkali lignin | H<sub>2</sub>O, EtOH, toluene (6:6:8), 7.5 g H<sub>2</sub>SO<sub>4</sub> |
| YM CL2E | Japanese cedar alkali lignin | H<sub>2</sub>O, EtOH, toluene (6:6:8), 7.5 g H<sub>2</sub>SO<sub>4</sub> |
| YM CL2A | Japanese cedar alkali lignin | H<sub>2</sub>O, EtOH, toluene (6:6:8), 7.5 g H<sub>2</sub>SO<sub>4</sub> |
| MY E1 | ball-milled *E. globulus* wood flour | AcOH containing 1% peracetic acid |
| MY E2 | ball-milled *E. globulus* wood flour | AcOH containing 1% peracetic acid |
| MY E3 | ball-milled *E. globulus* wood flour | AcOH containing 1% peracetic acid |
| MK EL1 | *E. globulus* alkali lignin | D<sub>2</sub>O, CD<sub>3</sub>COOD (2:1), 1.67 mM TsCl |
| MK EL2 | *E. globulus* cellulase-treated residual lignin | D<sub>2</sub>O, CD<sub>3</sub>COOD (2:1), 1.67 mM TsCl |

**Table 2.** MASL reaction details used in this study.

| Sample | volume (ml) | Reaction |  | Extraction solvent | Microwave reactor |
| --- | --- | --- | --- | --- | --- |
|  |  | temp (°C) | time (min) |  |  |
| YM C1T | 20 | 180 | 30 | toluene | Biotage Initiator<sup>+</sup> |
| YM C1E | 20 | 180 | 30 | ethyl acetate | Biotage Initiator<sup>+</sup> |
| YM C1A | 20 | 180 | 30 | acetone | Biotage Initiator<sup>+</sup> |
| YM E1T | 20 | 180 | 30 | toluene | Biotage Initiator<sup>+</sup> |
| YM E1E | 20 | 180 | 30 | ethyl acetate | Biotage Initiator<sup>+</sup> |
| YM E1A | 20 | 180 | 30 | acetone | Biotage Initiator<sup>+</sup> |
| YM C2T | 20 | 180 | 30 | toluene | Biotage Initiator<sup>+</sup> |
| YM C2E | 20 | 180 | 30 | ethyl acetate | Biotage Initiator<sup>+</sup> |
| YM C2A | 20 | 180 | 30 | acetone | Biotage Initiator<sup>+</sup> |
| YM E2T | 20 | 180 | 30 | toluene | Biotage Initiator<sup>+</sup> |
| YM E2E | 20 | 180 | 30 | ethyl acetate | Biotage Initiator<sup>+</sup> |
| YM E2A | 200 | 180 | 30 | acetone | Milestone StartSYNTH |
| YM CL1T | 200 | 180 | 30 | toluene | Milestone StartSYNTH |
| YM CL1E | 200 | 180 | 30 | ethyl acetate | Milestone StartSYNTH |
| YM CL1A | 200 | 180 | 30 | acetone | Milestone StartSYNTH |
| YM CL2T | 200 | 180 | 30 | toluene | Milestone StartSYNTH |
| YM CL2E | 200 | 180 | 30 | ethyl acetate | Milestone StartSYNTH |
| YM CL2A | 20 | 180 | 30 | acetone | Milestone StartSYNTH |
| MY E1 | 20 | 50 | 10 | ethyl acetate | Biotage Initiator<sup>+</sup> |
| MY E2 | 20 | 100 | 10 | ethyl acetate | Biotage Initiator<sup>+</sup> |
| MY E3 | 20 | 140 | 10 | ethyl acetate | Biotage Initiator<sup>+</sup> |
| MK EL1 | 15 | 160 | 30 | ethyl acetate | Biotage Initiator<sup>+</sup> |
| MK EL2 | 15 | 160 | 60 | ethyl acetate | Milestone StartSYNTH |

Ball-milled *E. globulus* wood particles were decomposed with a mixture (20 mL) of acetic acid containing 1% peracetic acid using a microwave reactor (Biotage Initiator Plus) at 50, 100 and 140° C for 10 min. After removal of the solvent, low-molecular-mass lignin was extracted with acetone (Entry 19-21). *E. globulus* alkaline lignin was degraded by microwave solvolysis using a mixture (15 mL) of deuterated acetic acid and D_2_O (2:1) containing 1 mM TsCl in a microwave reactor (Biotage Initiator Plus) at 160° C for 30 min (Entry 22). After reactions, MASL was extracted with ethyl acetate. Cellulase-treated residual lignin after acidolysis with maleic acid as described [18], was degraded by microwave solvolysis using a mixture (15 mL) of deuterated acetic acid and D_2_O (2:1) containing 1 mM TsCl in a microwave reactor (Milestone StartSYNTH) at 160° C for 60 min (Entry 23). After reactions, MASL was extracted with ethyl acetate. All MASL was diluted to a concentration of 200 mg/mL in DMSO (Nacalai Tesque) and stored at 4° C in the dark. Samples were filtered through 0.22-µm filters before use. For detailed information, described the (Table 1 and Table 2).

### 2.2 Cell lines and culture conditions

A Lewis lung carcinoma cell line, LLC, was obtained from RIKEN BRC (RCB0558). A human alveolar adenocarcinoma cell line, A549, and a human fibrosarcoma cell line, HT1080, were purchased from ATCC (Numbers CCL-185 and CCL-121). aHDFs (normal adult human dermal fibroblasts) were purchased from ScienCell Research Laboratories (cat no.2320). All cells were cultured in Dulbecco’s modified Eagle’s medium (DMEM; Nacalai Tesque) supplemented with 10% fetal bovine serum (FBS; Equitech-Bio), 100 U/mL penicillin, 100 µg/mL streptomycin, and 100 mM non-essential amino acids (complete DMEM) at 37 °C in 5% CO2/95% humidified air. Cells were trypsinized before passage.

### 2.3 Cell viability assay and Estimation of CC_50_ values

Cell viability was determined using a water-soluble tetrazolium salt assay. Cells were seeded into 96-well plate at a density of 1 x 10_3_ cells per well, and the next day, MASL were added to each well. After culturing for 12 or 24 h, 2-(2-methoxy-4-nitrophenyl)-3-(4-ni-trophenyl)-5-(2,4-disulfophenyl)-2H-tetrazolium monosodium salt (WST-8) solution (Nacalai Tesque) was added to the wells. After 1 h of incubation at 37 °C, culture supernatant was transferred to new 96-well plates, and the absorbance of each well was measured using a microplate reader (Emax; Molecular Devices). CC_50_ values were calculated as described [6].

### 2.4 Flowcytometric analysis

For the Annexin-V/propidium iodide assay, LLC, A549, HT1080 and HDFs were treated with MASL for 24 h. Concentrations of MASL were 0.05 and 0.1 mg/mL (for LLC), 0.1 and 0.2 mg/mL (for A549 and HT1080) and 0.05, 0.1 and 0.2 mg/mL (for HDFs). Cells were detached using AccutaseTM. After washing twice with washing buffer (PBS/2% FBS), cells were stained with Annexin V-FITC and PI solution (Nacalai Tesque) for 15 min at room temperature in the dark.

Following incubation, binding buffer was added, and cells were analyzed with a BD FACS Calibur cytometer (Becton Dickinson). For each sample, 10,000 gated events were recorded. For cell surface staining with anti-Fas and anti-FasL antibodies, LLC cells were treated with MASL at concentrations of 0.05 and 0.1 mg/mL for 24 hours. Cells were detached using AccutaseTM and incubated for 1 h at room temperature in the dark with PE-conjugated anti-FAS (CD95/APO-1) monoclonal antibody (15A7) (eBioscience™, DriveSan Diego, CA) or PE-conjugated anti-Fas ligand (CD178) monoclonal antibody (MFL3) (eBioscience™). Cells were washed twice with washing buffer and analyzed using a FACS Calibur cytometer as above.

### 2.5 Real-time RT-PCR

LLC cells were treated with MASL at concentrations of 0.05 and 0.1 mg/mL for 24 hours. Total RNA was extracted from cells using ISOGEN II (Nippon Gene), and reverse-transcribed using ReverTra Ace qPCR RT Master Mix (Toyobo). Real-time RT-PCR was carried out using Real-Time PCR Master Mix (KAPA Biosystems) and following probes and primers on a 7300 Real-Time PCR System (Applied Biosystems, CA). Primer/probes for mouse TNF-a gene were purchased from Applied Bioscience (Mm00443258_m1). To detect mouse β-actin gene mRNA, forward (5’-ACGGCCAGGTCATCACTATTG) and reverse (5’-TGGATGCCACAGGATTCCAT) primers and a probe ( ACGAGCGGTTCCGAT)were used. All values (average ± SD) were normalized with regard to the β-actin mRNA level in each sample as follows: Relative mRNA level = [(target gene mRNA level in sample)/(β-actin gene mRNA level in sample)]/[(target gene mRNA level in untreated control)/(β-actin gene mRNA level in untreated control)].

### 2.6 Western blotting

LLC, A549, HT1080 and HDFs were treated with MASL at concentrations of 0.05-0.2 mg/mL for 24 hours. Cells were washed twice with PBS and lysed in RIPA buffer (Nacalai Tesque) for 10 min at 4°C. Lysates were agitated at room temperature and incubated on ice for 30 min. All samples were centrifuged at 20,600 × g for 5 min and supernatants were collected. Protein concentrations of supernatants were determined using bicinchoninic acid (BCA) assays, and each sample was loaded at 5 µg/lane onto 8 or 10% gel (Bis-Tris Gel NuPAGE®). Proteins were separated by electrophoresis in 5% MOPS SDS Running Buffer (Thermo Fisher Scientific), followed by dry-blotting onto nitrocellulose iBlot®gel transfer stacks using iBlot Gel transfer device (Thermo Fisher Scientific) for 7 min. Membranes were washed in TBST (0.02 M Tris-HCl pH7.5, 0.15 M NaCl, 0.1% Tween 20), and blocked in Blocking One solution (Nacalai Tesque) for 1 h at room temperature. Membranes were incubated overnight at 4°C with following primary antibodies: anti-NF-κB p65, anti-phospho-NF-κB p65, anti-phosphor-mTOR (Ser2448), anti-mTOR (7C10) and mouse anti-human β-actin antibodies. All primary antibodies were purchased from Cell Signaling Technology and used at a dilution of 1:1,000, except that anti-β-actin antibody was purchased from Sigma-Aldrich (catalog no. A2228) and used at 1:4,000. Membranes were then washed three times with TBST and incubated with peroxidase-conjugated anti-mouse IgG secondary antibody (1:4,000 dilution) (Sigma-Aldrich; catalog no. A4416) at room temperature for 1 h. After washing three times with TBST, membranes were treated with enhanced chemiluminescence detection reagents (Chemi-Lumi One Super; Nacalai Tesque) and exposed to ECL Select LAS500 (GE Healthcare, Buckinghamshire, UK). All images were scanned with Image J software.

### 2.7 LLC tumor mouse model

Animal experiments were performed with approval from the Experimental Animals Committee, Kyoto Prefectural University of Medicine (Code No M29-137), and all procedures were followed in accordance with the NIH Guide for the Care and Use of Laboratory Animals. Six-week-old female C57BL/6 mice were purchased from Shimizu Laboratory Supplies (Kyoto, Japan). As a tumor model, LLC cells (1.0 × 10_5_) resuspended in 100 µL PBS were subcutaneously injected into the flanks of 7-weeks old mice. After tumors grew to a size of approximately 100 mm_3_, mice were given intraperitoneal injections of 0.1% MASL at a dose of 20 mg/kg body weight or DMSO as a control every other day for 14 days. At 2-day intervals, mice were weighed, while tumor size was measured with a caliper. Tumor volume was calculated as follows: tumor volume = d_2_ × D/2, where d and D were the shortest and longest diameters, respectively. After mice were euthanized, tumors were weighed and sera were tested for concentrations of alanine aminotransferase (AST) and aspartate aminotransferase (ALT) (Blood liver enzyme assay; Oriental Yeast, Osaka, Japan), 14 d after implantation of LLC. Body weight was checked at 2-day intervals.

### 2.8 Immunohistochemistry

LLC tumors were surgically removed from mice 14 days after initiation of YM CL1T treatment. Specimens were cryosectioned into 10-µm slices, followed by fixation with 2% (vol/vol) paraformaldehyde in PBS and blocking with 5% (vol/vol) normal goat serum, 1% BSA, and 0.3% Triton X-100 in PBS. Sections were incubated with primary antibodies rabbit polyclonal anti-caspase 3 and rat monoclonal anti-CD31/PECAM-1 antibody (Novus Biologicals, Centennial, CO) in 1% normal goat serum, 1% BSA, 0.3% Triton X-100 in PBS, before washing and incubation with secondary antibodies, Alexa Fluor 488-conjugated goat anti-rabbit and Alexa Fluor 568-conjugated anti-rat antibodies (Invitrogen). Cell nuclei were also stained with Hoechst 33342 Solution (Thermo Fisher Scientific). To quantify percentages of caspase 3-expressing areas, tumor sections were immunostained. At least 3 images per tumor were acquired (three tumors per group). As above, except that Cy3-conjugated goat anti-rabbit IgG antibody (Jackson Immuno-Research, West Grove, PA) was used as a secondary antibody. Acquired images were analyzed using Image J software (National Institutes of Health). Caspase 3-positive areas were also recognized with Image J software, and the total percentage of Caspase 3-positive areas for each image was calculated.

### 2.9 Statistical analysis

All experimental data are shown as the mean ± standard deviation (SD). Two-tailed Student’s t tests and parametric one-way or two-way analysis of variance (ANOVA) were used to analyze differences among groups. The Tukey-Kramer post-hoc test was applied to determine specific differences among the groups. In all analyses, P<0.05 was regarded as statistically significant. Statistical analysis was carried out using GraphPad Prism 6 (GraphPad Software, Inc).

## 3 Results

### 3.1 MASL treatments reduced viability of tumor cells, whereas normal cells were less remarkably affected

We prepared lignin samples from *E. globlus* and Japanese cedar wood through microwave acid-catalyzed solvolysis procedures (Table 1 and Table 2). Out of twenty-three samples of MASL tested for their impact on tumor cell viability, eight MASL samples significantly reduced the viability of LLC cells (Figure 1) (Table S1). Specifically, YM E2T, YM CL1T, YM CL2T and MKEL2 showed higher inhibitory efficacies, with the concentration-dependent manner evident among the twenty-three MASL samples (Figure 1A) (TableS1). Based on these results, we selected YM E2T, YM CL1T and YM CL2T samples for further experiments to analyze their anti-tumor efficacies in more detail. Due to limitations in the available MKEL2 sample, only in-vitro experiments were conducted. Three tumor cell lines of LLC, A549 and HT1080 were treated with MASL at 0.4, 0.2, 0.1 and 0.05 mg/mL for 12-h and 24-h. CC_50_ values of the MASL samples were calculated based on cell viabilities (TableS2, TablsS3). The average of CC_50_ values on LLC were lower (12-h: 0.17 ± 0.07 mg/mL and 24-h: 0.11± 0.03) than A549 and HT1080 (12-h: 0.26 ± 0.05 mg/mL and 24-h: 0.16± 0.03 and 12-h: 0.28 ± 0.11 mg/mL and 24-h: 0.16± 0.07) (Figure 1B) (Figure S1). The concentration of each applicable MASL were determined corresponding to the CC_50_ value and applied to three cancer cell lines for further research.

**Figure 1.**
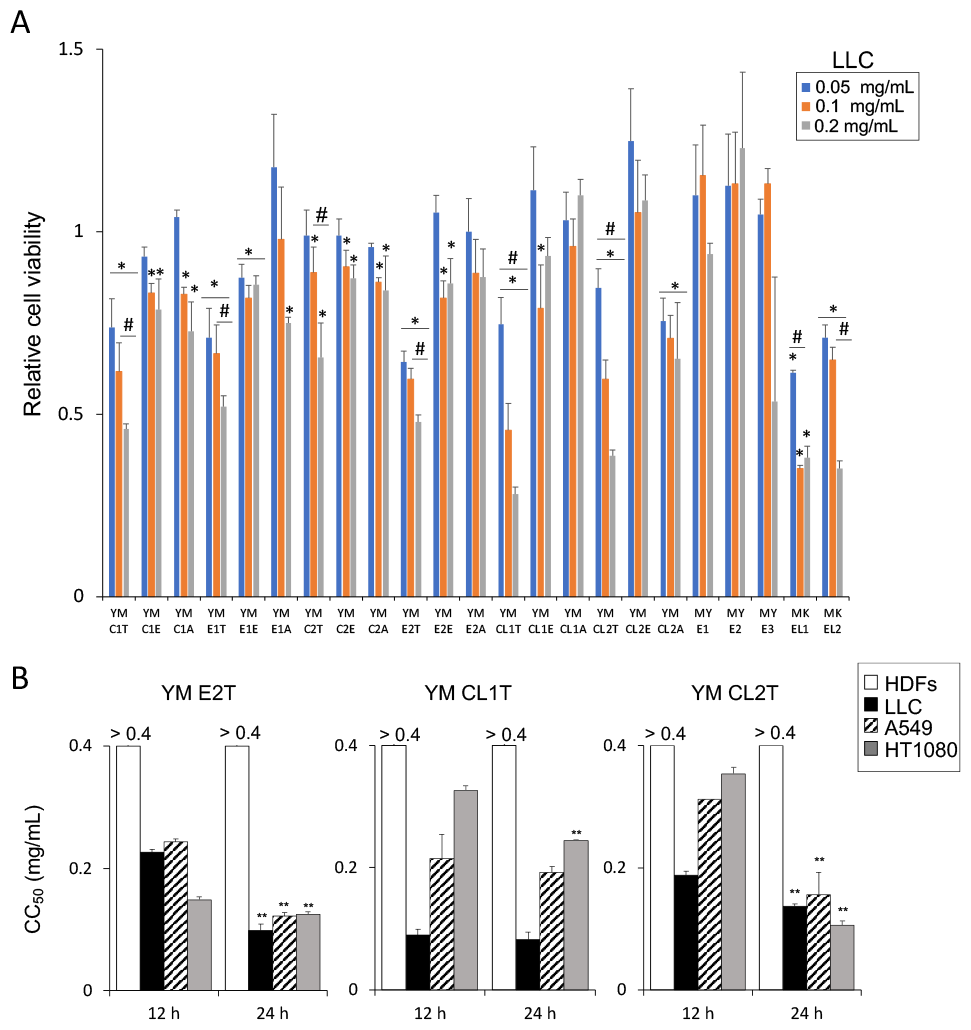
Effects of twenty-three MASL on tumor cell viability. (A) LLC were treated with 0.05, 0.1 and 0.2 mg/mL of the indicated MASL. Twenty-four h later, cell viability was determined using a water-soluble tetrazolium salt assay. Data are expressed as the mean ± SD (n=3). *p<0.05 vs. DMSO-treated control, and #p<0.05 between groups, by using repeated measures analysis of variance (ANOVA). (B) CC_50_ values of MASL on tumor cells and normal cells. The value of CC_50_ of MASL treatment on; LLC, A549 and HT1080 and HDFs were calculated based on the cell viability as described in Materials and Methods. Data are expressed as the mean ± SD (n=4). **p<0.01 vs. 12-h, by two-tailed Student’s t test.

### 3.2 Tumor cells underwent apoptosis after MASL treatment

According to the results of CC_50_ evaluation, we looked at the cell death status of tumor cells treated with MASL by Annexin V-FITC/PI staining (Figure 2). The concentration of each MASL samples were determined based on the CC_50_ values of each cell lines (Figure 1B). Following the CC_50_ values, we treated LLC cells with 0.05 and 0.1 mg/mL of MASL, while higher concentrations of MASL (0.1 and 0.2 mg/mL) were supplied to A549 and HT1080 cell treatments. HDFs were treated with 0.05, 0.1 and 0.2 mg/mL of MASL as control. Thus, we were able to directly compare the cell death condition of each tumor cell line with those of HDFs. As a result, percentages of early- and late-stage apoptotic cells (Annexin V + PI - and Annexin V + PI +, respectively) were increased in a dose-dependent manner by treatment with MASL (Figures 2, S2 and S3). In sharp contrast, there was no evidence of apoptosis induction in HDFs treated with MASL except for 0.2 mg/mL of YM E2T (Figure 2). These findings strongly suggest pro-apoptotic effects were observed in MASL treatments on tumor cells.

**Figure 2.**
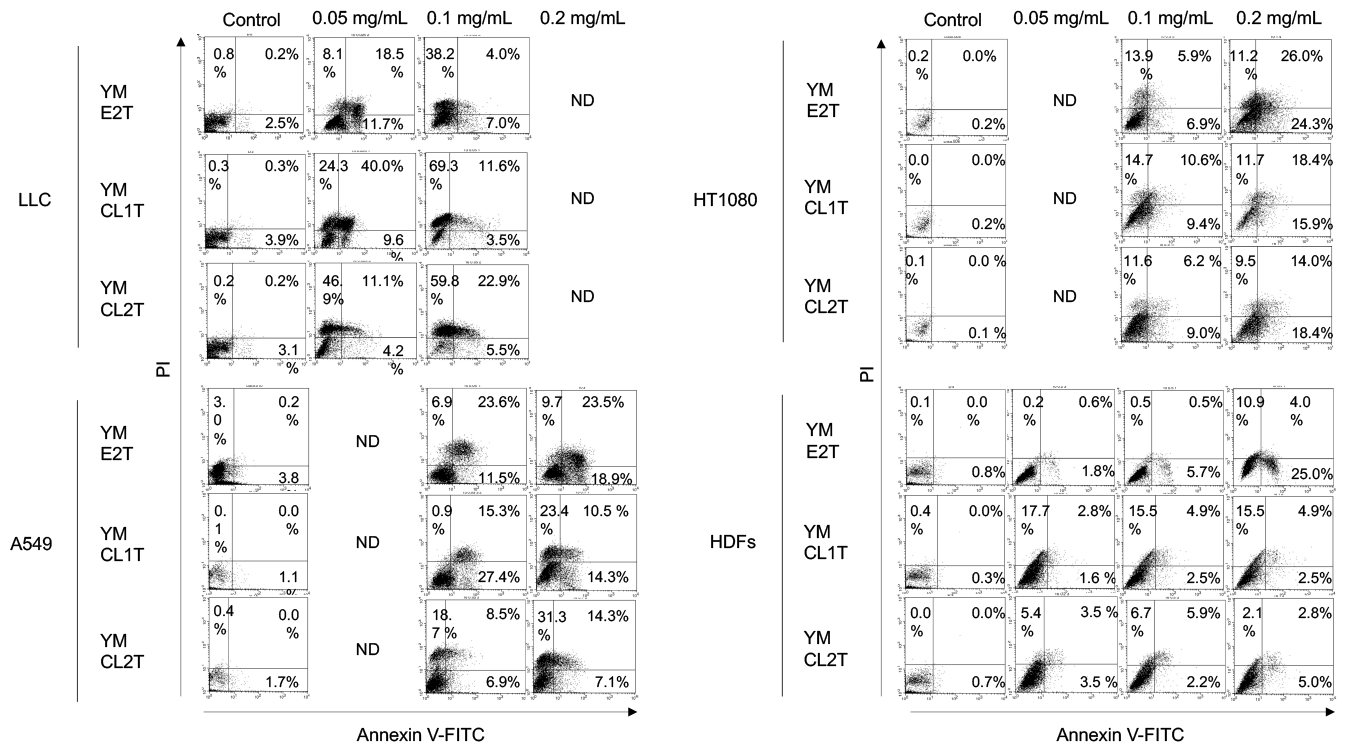
MASL treatment triggered apoptosis in tumor cells. LLC, A549, HT1080 and HDFs were treated with the indicated MASL for 24 hours, followed by Annexin V-FITC/PI staining and flowcytometric analysis. Dot plots of the representative samples are shown. N.D., not determined.

### 3.3 MASL treatment modulated apoptotic pathways in tumor cells

To understand molecular mechanisms of apoptosis induction by MASL treatments in tumor cells, we examined whether MASL activated expression levels of mRNA and protein for TNF-α, Fas, and FasL that could contribute to extrinsic pathways of apoptosis induction. Real-time RT-PCR analysis revealed significant upregulation of TNF-α mRNA in the LLC cells treated with MASL (Figure 3A). MASL-treated LLC cells also exhibited increased protein expression levels of both Fas and FasL on the cell surface, as evidenced by flowcytometric analysis (Figure 3B). We additionally checked modulation mechanisms in intrinsic pathway at the protein levels. To clarify this, Western blot analysis was performed to estimate phosphorylation statuses of NF-κB p65 and mTOR.

**Figure 3.**
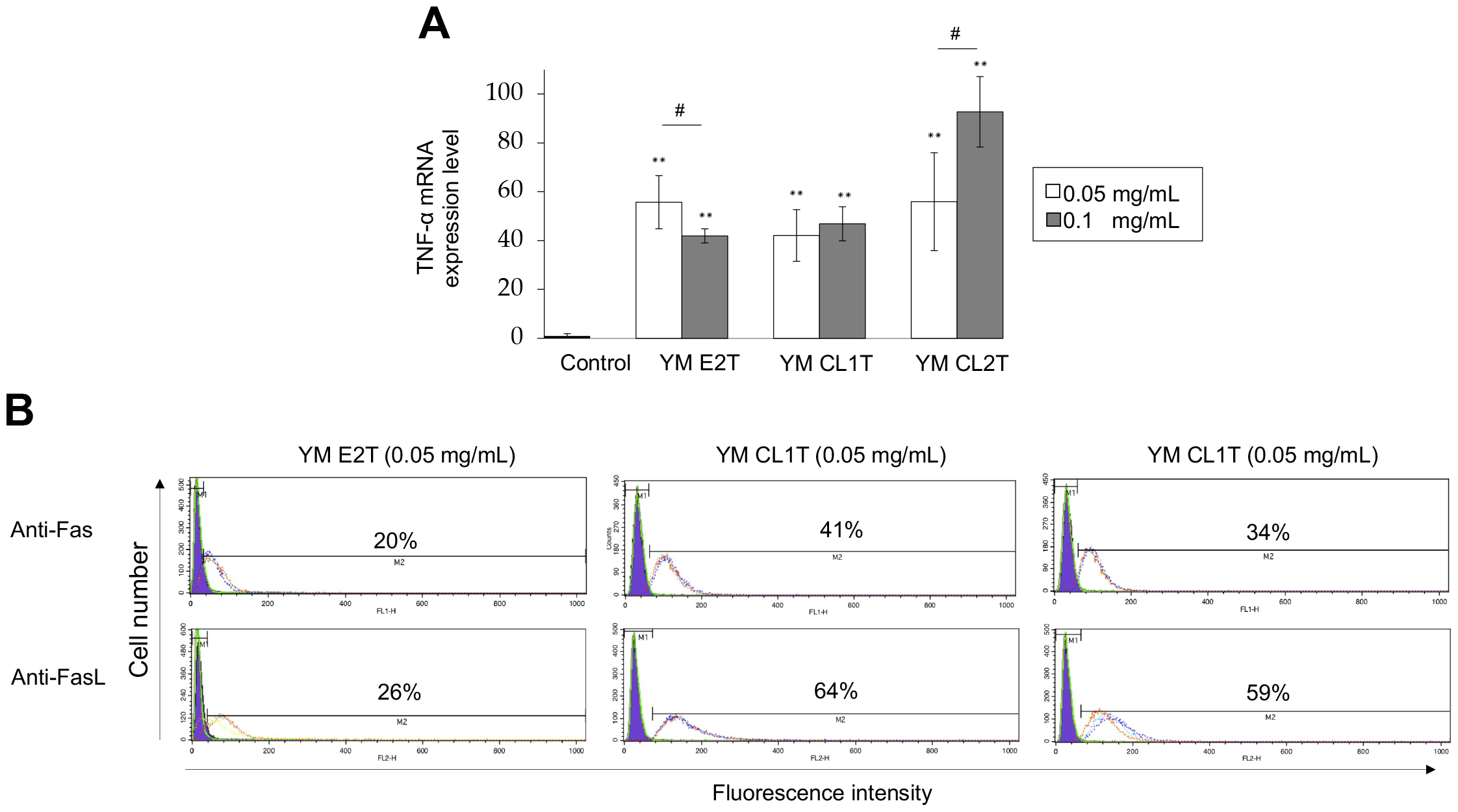
MASL-treated tumor cells expressed TNF-α, Fas and FasL. LLC cells were treated with the indicated MASL for 24 h. (A) RNA was obtained from the cells, and mRNA levels for the TNF-α gene were analyzed by real-time-RT-PCR. Data are expressed as the mean ± SD (n = 3). **P<0.01 vs. DMSO-treated control; # p<0.05, between groups, by Tukey-Kramer test. (B) Cells were stained with PE-conjugated anti-Fas and anti-FasL antibodies and analyzed by flowcytometry. Histograms for DMSO-treated cells (controls, purple) and MASL-treated cells (transparent) are shown. Three independent cell aliquots were tested for each group and shown with red, blue, and green lines, respectively. Percentages represent average proportions of positive cells.

Intriguingly, MASL treatment remarkably reduced phosphorylated p65 in LLC, A549, and HT108 cells. Phosphorylation of mTOR was suppressed at various degrees in the tumor cells cultured with MASL (Figure 4). These results strongly suggest that MASL modulated various extracellular and intracellular anti-apoptotic pathways in tumor cells to induce apoptosis.

**Figure 4.**
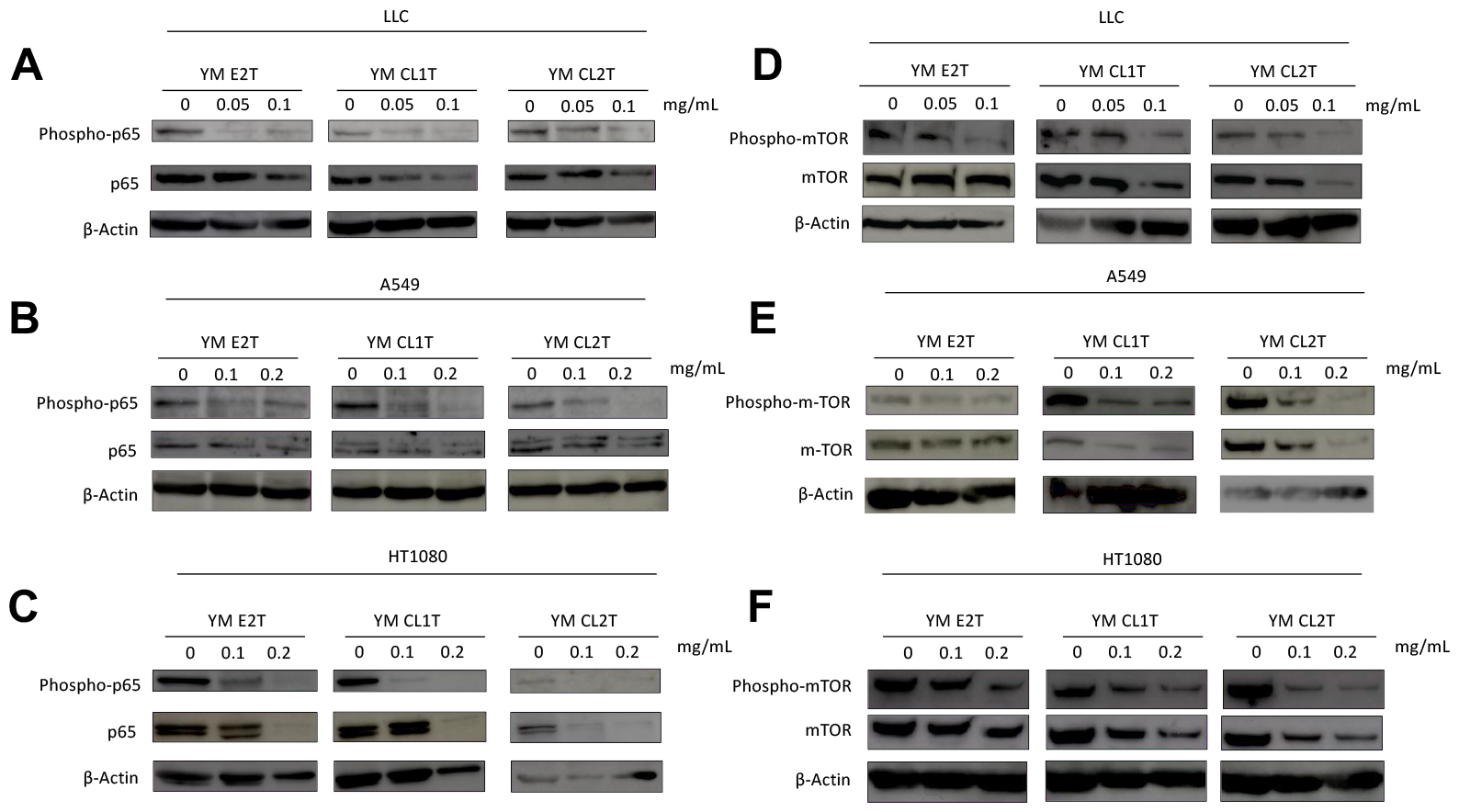
MASL inhibited NF-κB signaling, while modifying intracellular anti-apoptotic signaling in cancer cells. LLC, A549 and HT1080 cells were treated with the indicated MASL for 24 h. Cells were extracted and subjected to western blot analysis. (A, B and C) Anti-phospho-NF-κB p65, anti-NF-κB p65, and anti-β-actin antibodies were used. (D, E and F) Anti-phospho-mTOR (Ser2448), anti-mTOR and anti-human β-actin antibodies were used.

### 3.5 YM CL1T administration inhibited tumor growth in vivo

To access the potential anti-tumor effects of MASL in vivo, we establish a Lewis lung carcinoma mouse. Tumor-bearing mice were intraperitoneally administrated YM E2T, YM CL1T and YM CL2T. The YM CL1T significantly inhibited increase in tumor volume and weight (Figure 5A&B). In contrast to YM CL1T, although the average of the tumor volume was less than control in YM E2T and YM CL2, we did not observe significant differences in this research. Meanwhile, MASL-treatment caused neither significant body weight loss (Figure 5C) nor liver toxicity in the tumor-bearer (Figure 5D). These results suggest that YM CL1T has significant anti-tumor effects in vivo in mice, without causing severe adverse events.

**Figure 5.**
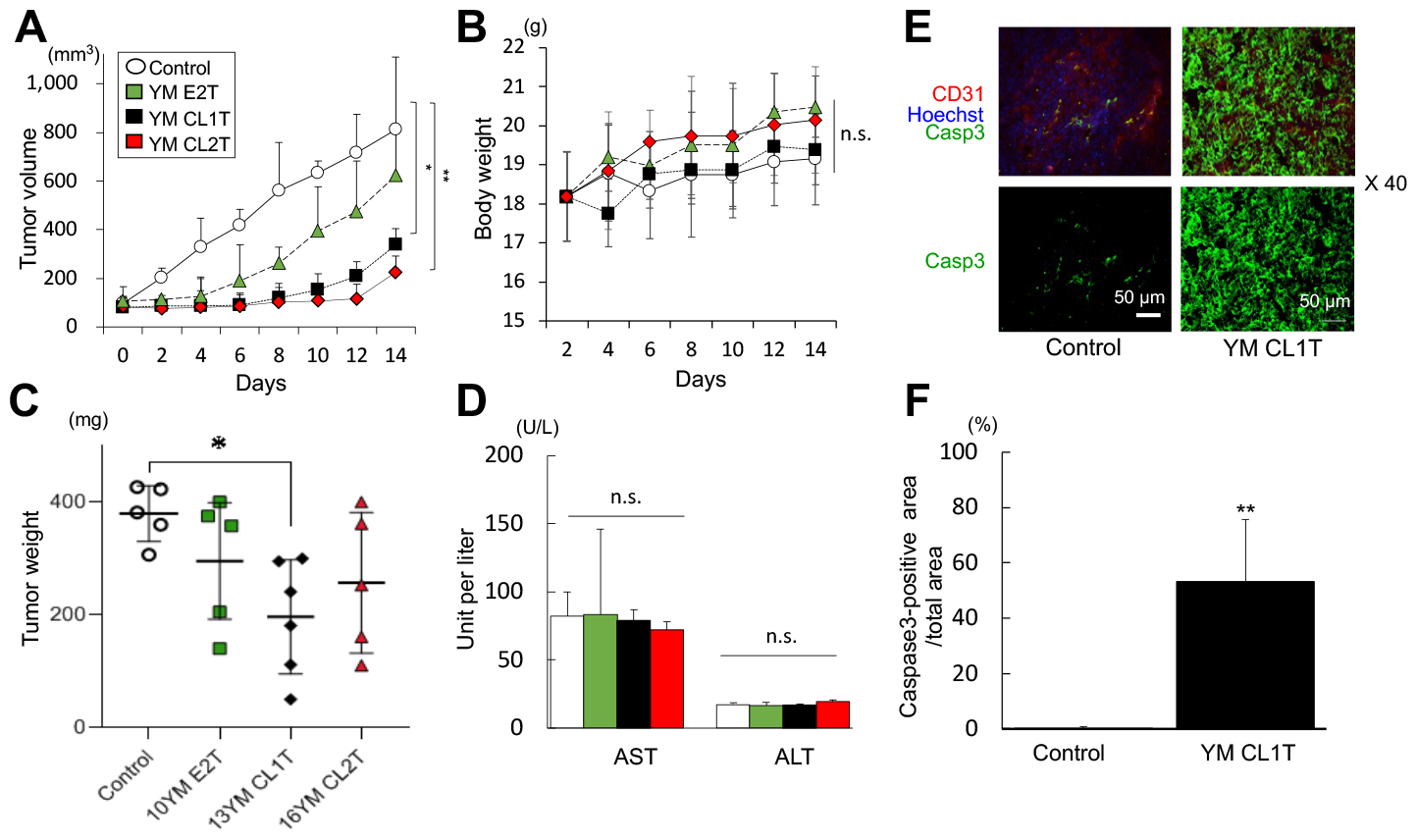
Administration of YM CL1T significantly inhibited growth of LLC tumors in mice. LLC tumor-bearing mice were given intraperitoneal injections of the indicated MASL. Control mice were injected with DMSO. Mice were euthanized 14 days after initiation of MASL treatment. (A) Kinetic changes in tumor sizes (mean ± SD) are plotted. (B) Weights of tumors on Day 14 are shown. MASL was injected intraperitoneally at a dosage of 20 mg/kg body weight. (C) Kinetic changes in body weights of mice are shown. (D) Sera were collected from the mice on day 14 and tested for concentrations of AST and ALT. Data are expressed as mean ± SD. N=5 mice (for Control, YM E2T and YM CL2T groups) or 6 mice (YM CL1T group). (E) LLC tumor specimens were subjected to immunohistochemical analysis using anti-caspase 3 and CD31 antibodies and Hoechst 33342. Representative fluorescent microscopic images of tumor sections are shown. Original optical magnification was x 40. Size of the scale bar = 50 µm in length. (F) Caspase 3-positive areas (mean ± SEM) were calculated from fluorescent microscopic images, and ratios of caspase 3-positive area per total area are shown. In (b), data are expressed as means ± SD. n = 3 per tumor section. Three tumors for each group. *p<0.05 and **p<0.01 between groups, by ANOAVA. N.S., not significant.

3.6 YM CL1T suppressed tumor growth in association with elevated expression of caspase 3 and inhibition of angiogenesis in tumors.

Finally, to clarify mechanisms underlying the anti-tumor activity of YM CL1T in vivo, we assessed expression of caspase 3 in tumor tissue two weeks after YM CL1T treatment. Caspase 3 was more strongly and broadly expressed in tumor tissues from the mice treated with YM CL1T compared with tumor tissues from DMSO-treated control mice (Figure 5E, F). Statistical evaluation confirmed a significant expression of caspase 3 effect of YM CL1T (Figure 5F). These results strongly suggest that YM CL1T promoted apoptosis induction in tumor cells in vivo.

## 4. Discussion

The physiological functions of lignin and lignin-containing fractions from plant cell walls have been extensively studied [5, 19-22]. However, the molecular mechanisms of their tumor suppression effects in-vivo and in-vitro have not been fully explained [3]. In this study, we prepared twenty-three MASL fractions from Japanese cedar and *E. globulus* wood to investigate the anti-tumor properties of MASL in cells and mice. We observed a reduction in cell survival, due to apoptotic cell death in tumor cell lines when treated with YM E2T, YM CL1T and YM CL2T from MASL. To elucidate the cell death pathway of the MASL-mediated anti-tumor effects, we focused on the main pro-apoptotic and anti-apoptotic pathways in tumor cells [23]. From the FACS and real-time RT-PCR analysis, we have revealed that the extrinsic pathway (Death Receptor Pathway) contributes to MASL-triggered cell death. We have confirmed the upregulation of Fas (CD95/APO-1), FasL, and TNF-α (Figure 3). Previous studies have reported that cellulase-treated lignin-carbohydrate complexes (LCCs) activated myeloid dendritic cells via Toll-like receptor 4 (TLR4) [5]. Apoptosis caused by the signaling through the TNF-receptor superfamily and other types of death receptors n is widely known [24]. The regulation of TNF alpha and TRAIL receptor is preferable to be analyzed in future research, treating with MASL. In addition to the extrinsic pathway, apoptotic stimuli, including Bax, are involved in a mitochondria-mediated apoptotic pathway, known as the intrinsic pathway, in tumor cells [25]. During this research, we were unable to confirm the differentiation in Bax and Bcl-2 protein levels. Therefore, it is suggested that the MASL-induced cell death is related with extrinsic pathway than intrinsic pathway of apoptosis. The western-blot analysis further suggested the contribution of NF-κB pathway and MAPK pathway modulation to the tumoricidal effects of MASL; mTOR has been shown to suppress NF-κB activity (Figure 4). The reduction of mTOR can decrease the expression of anti-apoptotic genes regulated by NF-κB, thereby contributing to cell death [26]. Furthermore, confirming the interaction between the MAPK pathway, mTOR, and ERK will provide insight into prospective cell survival and proliferation mechanisms induced by MASL treatments [27].

Finally, we revealed that the YM CL2T-treated mouse group exhibited remarkable anti-tumor effects, including the suppression of tumor growth and upregulation of caspase 3, while other groups did not. This research has provided us with two hypotheses. First, the MASL killed tumor cells more effectively in in-vitro than in in-vivo. Secondly, although in this research we focused only on the apoptotic pathway—programmed cell death—modulated by MASL treatments, it may also be related to uncontrolled cell death, such as necrosis. The role of necroptosis in tumorigenesis is still not fully understood, as recent studies have reported both tumor-promoting and tumor-suppressing effects of necroptosis, and this aspect can be applied in this research [28].

Taken together, MASL induced cell death in mammalian tumor cells by modulating the apoptotic pathways. We obtained signals related to tumor cell death for MASL treatments, allowing us to perform the first in-vivo study of MASL treatments in the tumor mouse model with the sufficient information on MASL reagents’ reactions. Our results indicate that MASL has potential anti-tumor effects. We hope that our data will be useful as a platform for future experiments on lignin-related compounds and cell death mechanisms. This molecular understanding will also serve as an important dataset for future drug development projects. Our data opens up promising avenues of research to better understand this fascinating example of sustainable resource application in medicine.

## SUMMARY

The analysis of MASL showed remarkable tumor suppression both in vivo and in vitro experiments. Apoptosis of tumor cells was associated with augmented expression of TNF-α, Fas and FasL, as well as suppression of the antiapoptotic NF-κB and mTOR pathways. Repetitive intra-peritoneal administrations of YM CL1T significantly suppressed growth of LLC tumors in mice with elevated caspase 3 expression in the tumor tissue. Therefore, MASL may have potential as a novel anti-tumor agent, as suggested by our interdisciplinary study between biomedical and biomass research fields.

## Supplementary Materials

The following are available online at www.mdpi.com/xxx/s1, Figure S1: CC_50_ values of MK EL2 on tumor cells and normal cells; Figure S2. Apoptotic cells were more numerous in tumor cells treated with MASL; Figure S3: Apoptotic death of tumor cells induced by MK EL2.

## Author Contributions

Conceptualization, RK, OM, WH; methodology, RK, EO, YM, TM, CK, MS, HN, GP writing—original draft preparation, RK, YM, OM, WH. All authors have read and agreed to the published version of the manuscript.

## Funding

Part of this work was supported by the New Energy and Industrial Technology Development Organization (NEDO), the Japan Science and Technology Agency (JST), Analysis and Development System for Advanced Materials (ADAM) and Mission 5-1 Grant of RISH, Kyoto University. This research was supporting by Department of Immunology, Graduate School of Medical Scienceand Laboratory of Biomass Conversion, Research Institute for Sustainable Humanosphere (RISH), Kyoto University.

## Acknowledgments

We gratefully acknowledge the New Energy and Industrial Technology Development Organization (NEDO), the Japan Science and Technology Agency (JST), Analysis and Development System for Advanced Materials (ADAM) and Mission 5-1 Grant of RISH, Kyoto University and Dr. Steven D. Aird for editing of the manuscript and helpful comments.

## Conflicts of Interest

The authors declare no conflict of interest.

## Notes

### Competing Interest Statement

The authors have declared no competing interest.

