## Supplemental Figure 1-3 for "Lignin isolated by microwave-assisted acid-catalyzed solvolysis induced cell death on mammalian tumor cells by modulating apoptotic pathways"

Figure S1

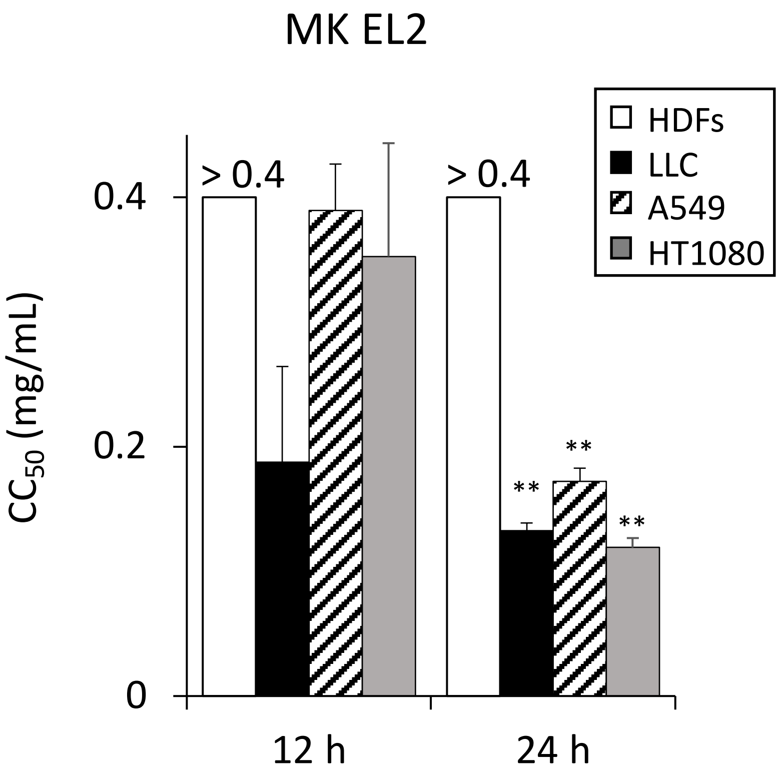


**Figure S1: CC50 values of MK EL2 on tumor cells and normal cells.**
The value of CC50 of MASL treatment on; LLC, A549 and HT1080 and HDFs were calculated based on the cell viability as described in Materials and Methods. Data are expressed as the mean ± SD (n=4). **p<0.01 vs. 12-h, by two-tailed Student's t test.

Figure S2

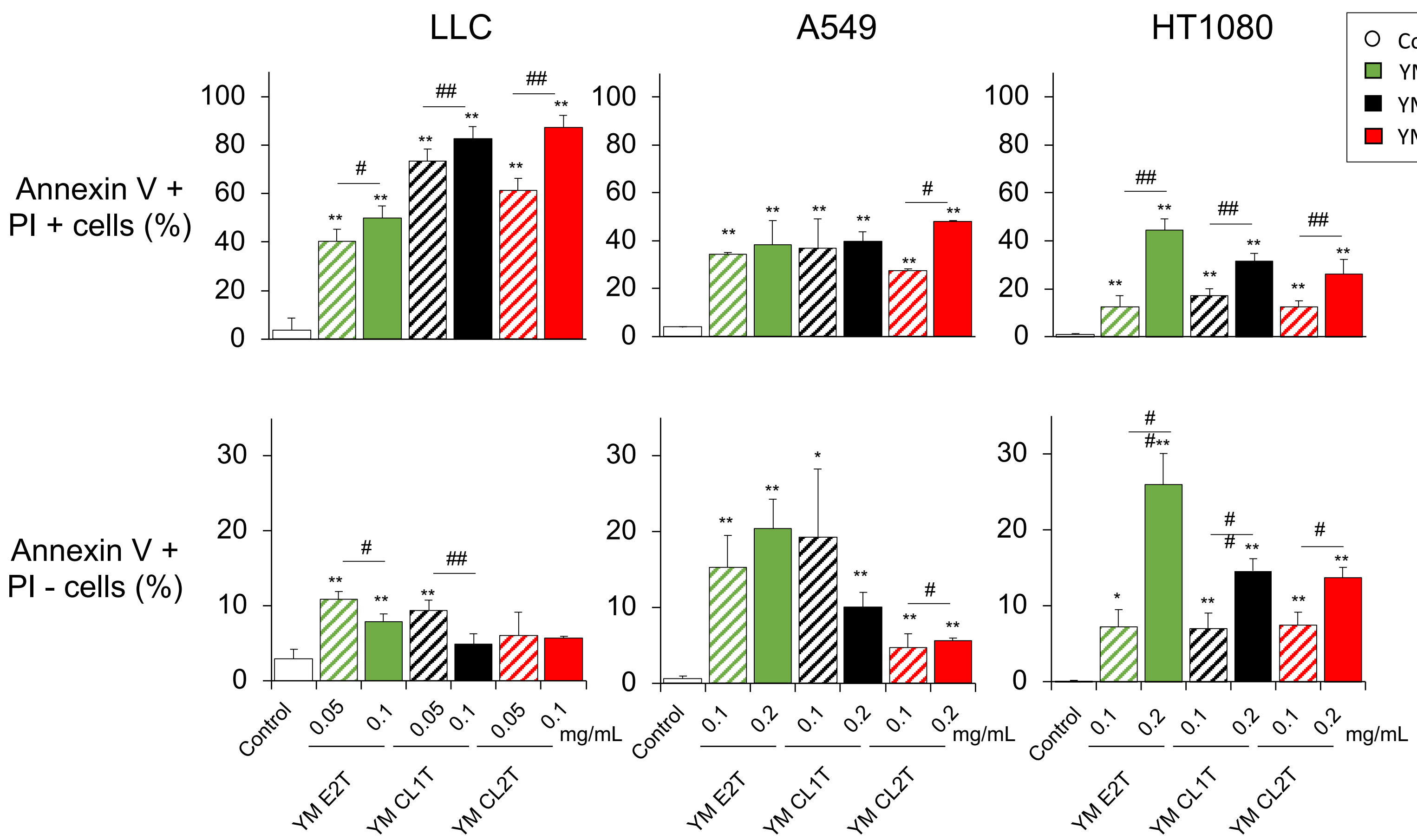


**Figure S2: Apoptotic cells were more numerous in tumor cells treated with MASL.** Percentages of Annexin V+ PI - and Annexin V + PI + cells were calculated based on the data shown in Fig. 4. Solid and stripe color shown for high and lower concentration of MASL treatments on each cell LLC; 0.05 and 0.1 mg/mL, A549 & HT1080; 0.1 and 0.2 mg/mL. Data are expressed as the mean ± SD (n=3). *p<0.05 and **p<0.01, vs. control; #p<0.05 and ##p<0.01, between groups, by Turkey-Kramer test.

Figure S3

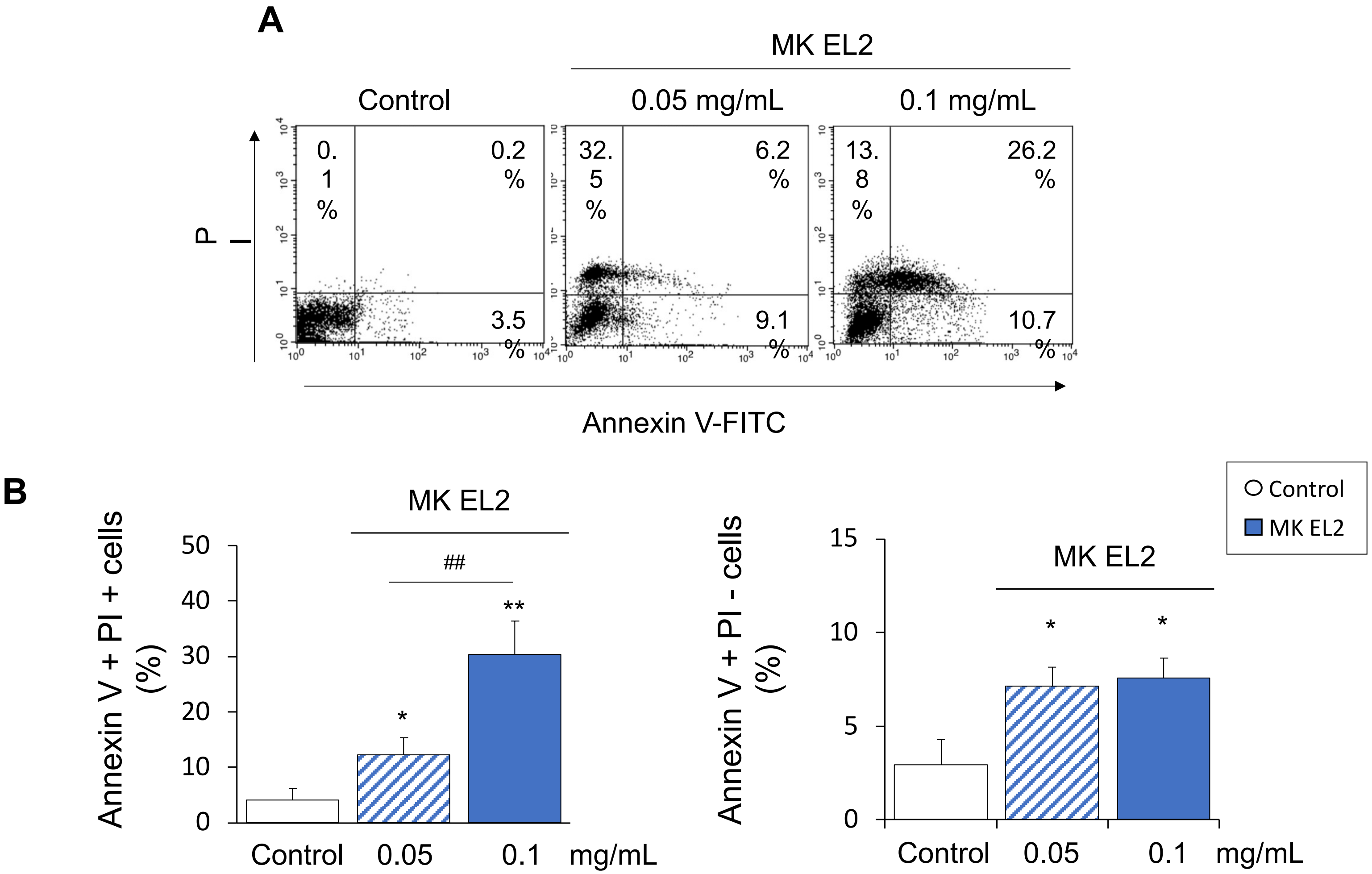


**Figure S3: Apoptotic death of tumor cells induced by MK EL2.**
LLC cells were treated with the indicated concentrations of MK EL2 for 24 h. Apoptosis was estimated by flowcytometry following Annexin V-FITC and PI staining. (A) Dot plots of the representative samples are shown. (B) Percentages of Annexin V + PI + and Annexin V + PI - cells were calculated based on the data shown in (A). Solid and stripe color shown for high and lower concentration of MASL treatments on each cell LLC; 0.05 and 0.1 mg/mL. Data are expressed as the mean ± SD (n=4). *p<0.05 and **p<0.01, vs. control; # p<0.05 and ##p<0.01, between groups, by Turkey-Kramer test.
